# *Staphylococcus aureus*-Induced Degeneration of Nociceptive Neurons in *Caenorhabditis elegans*

**DOI:** 10.1101/2025.05.01.651706

**Authors:** Elizabeth M. DiLoreto, Xavier Gonzalez, Khursheed A. Wani, Jiali Shen, Javier E. Irazoqui, Jagan Srinivasan

## Abstract

**Background:** In all animals, the nervous system senses microbial signals to influence host defense. Despite emerging as important sensors of infection to regulate immunity and inflammation, the mechanisms by which pain-sensing nociceptor neurons can detect infections are poorly defined. Using *C. elegans* as a tractable model host that shares many features with mammalian systems, we investigated nociceptor function during bacterial infection.

**Results:** *In vivo* intracellular Ca^2+^ imaging of nociceptor ASH neurons revealed a drastic reduction in ASH responses to aversive stimuli in *Staphylococcus aureus-*infected animals compared to noninfected controls. Morphological examination showed that the ASH neurons lost integrity in the sensory processes that extend to the mouth, in a pathogen growth phase-dependent manner. Neighboring neurons did not exhibit this pathogen-induced neurodegeneration (PaIN) phenotype. Genetic analysis suggested that apoptosis, necrosis, ferroptosis, and autophagy are dispensable for the PaIN phenotype. In contrast, loss of the evolutionarily conserved stress-response transcription factor HLH-30/TFEB reduced the penetrance of ASH PaIN by about 50%. Moreover, infected animals showed defective ASH-mediated evasive behaviors, suggesting that the *S. aureus-* triggered drop in ASH activation and morphological degeneration are physiologically relevant.

**Conclusions:** Collectively, these findings reveal that nociceptor neurons lose functional and morphological integrity during infection with *S. aureus,* with severe consequences for animal behavior. Because *S. aureus* is a critical human pathogen, the induction of nociceptor PaIN may have important implications for human health.

## Background

Host-pathogen interactions are complex biological processes that engage multiple systems within an organism. Emerging evidence highlights the critical role of the nervous system in pathogen detection and immune regulation (1). Nociceptor neurons, specialized sensory neurons that detect noxious stimuli, have been implicated in sensing various pathogens and modulating inflammatory responses during infection (2). Nociceptive neurons have also been activated in the response to urinary tract infections caused by *Escherichia coli* in mice (3). Despite these advances in understanding how the nervous system orchestrates host responses and inflammation through nociceptor neurons, how pathogens influence the function of nociceptors remain poorly understood.

The genetic tractability of *Caenorhabditis elegans*, combined with its transparent body and simple nervous system, allows for precise manipulation and real-time visualization of neuronal activity during host-pathogen interactions, making it powerful model for elucidating neuro-immune communication. Many mechanisms of *C. elegans* neuronal function are conserved in mammals (4). Research using *C. elegans* has illuminated nervous system-mediated host defense through behavioral and molecular strategies. Behavioral defense encompasses pathogen avoidance strategies, where sensory neurons detect microbial cues and trigger evasive locomotion patterns to minimize pathogen exposure (5). These neural responses as a result of bacterial infection have also exhibited pathogen and neuron specific impacts (6). Molecular defense involves the activation of innate immune host defense genes, including the production of antimicrobial peptides that directly combat pathogens in the intestinal epithelium, and cytoprotective factors that repair the cellular damage caused by infection (7–11). Researchers have found that amphid neurons, located in the head of *C. elegans*, play a role in both behavioral and molecular host defense responses (7, 12).

We previously showed that the muscarinic-Wnt signaling cascade is a key pathway in *C. elegans* host defense against infection (8). Upon intestinal colonization by the human pathogenic bacterium *Staphylococcus aureus*, *C. elegans* exhibits increased acetylcholine signaling, which activates the Wnt signaling pathway in intestinal epithelial cells through muscarinic acetylcholine receptors (8). Wnt signaling leads to the production of antimicrobial peptides such as CLEC-60, a C-type lectin that is functionally analogous to mammalian antimicrobial C-type lectins (8, 13). In *C. elegans,* only neurons synthesize and release acetylcholine, implicating the nervous system in the host response during *S. aureus* infection (14). Although the downstream components of the cholinergic Wnt pathway are better understood, the upstream neuronal circuits that trigger the initial cholinergic response remain unidentified. Understanding these upstream neuronal pathways is crucial for elucidating how the nervous system integrates pathogen detection with the activation of appropriate behavioral and molecular host defense mechanisms and may provide insights into conserved neuro-immune interactions across species.

ASH (Amphid Single Cilium H, left/right pair) neurons in *C. elegans* are polymodal nociceptors that respond to a diverse array of aversive stimuli, including strong mechanical stress, osmotic shock, chemical repellents, and pH changes (15–19). These bilaterally symmetric sensory neurons project dendrites to openings near the mouth, allowing direct contact with the external environment, and have cell bodies positioned in the head near the nerve ring (15, 20). ASH neurons have been shown to have distinct roles in mediating pathogen innate immunity during bacterial infection (5). ASH-ablated animals have increased innate host defense gene expression and increased survival during *Pseudomonas aeruginosa* infection, suggesting that ASH normally repress innate immune expression; however, the bacterial-sensing mechanisms have not been elucidated. Additional research has shown that in response to different pathogens, sensory neurons can each play a different role in sensation and survival (6). Forced ASH depolarization has been shown to evoke acetylcholine (ACh) release, suggesting that ASH might be able to sense *S. aureus* infection and trigger ACh release for Wnt pathway activation (8). (21).

In this study, we investigated the role of ASH neurons in *C. elegans* behavioral host defense against *S. aureus* infection by combining calcium imaging, genetic ablation, and behavioral assays. We demonstrate that infection leads to a rapid inhibition of ASH neuronal activity within 3 hours, followed by progressive neurodegeneration characterized by structural abnormalities in sensory dendrites. This pathogen-induced neurodegeneration (PaIN) is specific to ASH neurons and not observed in neighboring sensory neurons under identical conditions. Furthermore, we show that PaIN is independent of major cell death pathways but requires the transcription factor HLH-30/TFEB. HLH-30/TFEB is a master regulator of autophagy and lysosomal biogenesis that is also critical for the induction of molecular host defense in *C. elegans* and mammals (22). Thus, our findings suggest that HLH-30/TFEB plays an important role during *S. aureus* infection in ASH neurodegeneration. Furthermore, we find that infection impairs ASH-dependent aversive behaviors, suggesting that ASH inhibition and PaIN by *S. aureus* have important physiological consequences. These findings provide important insights into how bacterial pathogens can modify a host’s behavior by inducing the degeneration of nociceptor neuron form and function.

## Materials and Methods

### Strains, media, and culture conditions

*C. elegans* strains were maintained on 60 mm petri dishes containing 10 mL of nematode-growth media (NGM: 51 mM NaCl, 10 mM peptone, 51 mM agar, 1 mM CaCl_2_, 1 mM MgSO_4_, 25 mM KPO_4_ (pH 6.0), 13 µM Cholesterol) plates seeded with *E. coli* OP50 at 15 – 20°C, according to standard procedures (23). Strains used are listed in **Supplementary Table 1.** *S. aureus* SH1000 was grown normoxically in TSB (Sigma-Aldrich T8907) containing 50 µg/mL ampicillin for 16 hours at 37 °C with 150-200 RPM shaking in a beveled flask. 10 µL of culture was then plated on 35 mm petri dish containing 5 mL of Tryptic Soy Agar (TSA, BD 236950) and 10 µg/mL kanamycin and grown to confluency (37 °C for 6-8 hours, then 25 °C for 16 hours for dilute culture or 25 °C for 16 hours). Plates are stored at 4 °C until use and kept for 7 days. To grow *S. aureus* hypoxically, cultures were grown in minimally aerated (half volume of container filled with liquid) 15 mL conical tube, shaking at 150 RPM for 16 hours. For experiments, *E. coli* OP50 was grown shaking in beveled flask for 16 hours at 150 −200 RPM at 37 °C. 10 µL of *E. coli* was then plated on 35 mm NGM plates and incubated at 20 °C for 16 hours or 10 µL of *E. coli* was plated on 35 mm TSA plates without kanamycin and grown like *S. aureus* plates (at 37 °C for 6 hours, followed by 25 °C for 16 hours).

CeMbio cultures were grown under normoxic conditions at 25 °C with 170-200 RPM shaking in TSB (Sigma-Aldrich T8907) containing 50 µg/mL ampicillin for 20-24 h. 10 μL of the CeMbio culture was then spread on TSA (BD 236950) containing 50 µg/mL ampicillin at 25 °C for 20-24 hours. TSA containing 50 µg/mL ampicillin plates were used within 10 days. Ampicillin was included to prevent contamination by bacterial carry-over with the animals.

### Calcium Imaging

Imaging of neurons was performed as previously described (24, 25). ASH neurons were visualized using the CX10979 (N2;kyEx2865 [*sra-6p*::GCaMP3 @ 100 ng/µL]) *C. elegans* strain, generously provided by the lab of Cori Bargmann, PhD, which utilizes a genetically encoded calcium indicator green fluorescent protein-calmodulin-M13 peptide version 3 (GCaMP3) expressed in ASH neurons, ASH::GCaMP3 (26). Transgenic animals expressing the genetically encoded calcium sensor were picked as L4 stage animals. For experiments where the animals were infected prior to imaging, the animals were exposed to *S. aureus* grown on 35 mm Tryptic Soy Agar (TSA, BD 236950) and kanamycin 10 µg/mL plates or *E. coli* grown on 35 mm TSA plates for 3 hours at 25 °C prior to imaging. Animals were then immobilized in a microfluidic device and exposed to a buffer control (Pre), then the stimulus (Stimulus), and the stimulus was then removed and the buffer returned (Post). Preparation of the microfluidic device is described in (24). ASH is a light sensitive neuron (16). Animals were first photobleached using the blue fluorescent light until the neuron no longer responded to light, no more than 2 minutes of photobleaching. Animals were imaged for a 30 second trial; 5 second recording prior to stimulation, 10 second pulse of 1 M glycerol with 1 mM tetramisole in S. Basal, followed by a 15 second recording post stimulation. For the prolonged exposure experiments, it was a 150 second trial; 30 second recording prior to stimulation, 60 second pulse of *S. aureus* stimuli in TSB, followed by a 60 second recording post stimulation. The solutions used in imaging are all made in S. Basal medium (100 mM NaCl, 5.7 mM K_2_HPO_4_, 44 mM KH_2_PO_4_, 13 µM cholesterol in H_2_O). The solvent control solution was 1 mM tetramisole, 0.3 µM fluorescein in S. Basal. The flow control solution, not exposed to the animal but controlling the movement of the solutions in the olfactory chip was 1 mM tetramisole and 0.6 µM fluorescein in S. Basal. For the experiments in **Figure 3**, the flow control and solvent control are prepared in TSB, not S basal buffer.

Calcium imaging was performed on a Zeiss AOM Zeiss Axio Observer Microscope with H-F4-V2, Hamamatsu Flash 4.0 v2 Scientific CMOS camera, and X-120m, Excelitas 120 mini LED direct coupling light source. The initial sample alignment was performed with a Zeiss EC Plan-Neofluar 10x/0.3 M27 objective and image acquisition was performed with a Zeiss LD C-Apochromat 40x/1.1 UV-VIS-IR water immersion objective. Images were collected in V-software, VisiView software for image acquisition and ImageJ software for image analysis. Images were collected at one plane of view at a rate of 10 frames/second.

The soma of the ASH neuron in view was traced as the region of interest for fluorescence measurements. The fluorescence of an equally sized region was captured from the background. The fluorescence of the background was then subtracted from the neuron for each frame (equal to 0.1 seconds of the trial) to obtain the ΔF. The ΔF was then adjusted by dividing each frame by F_0_; this was calculated by the average ΔF from seconds 2-3 (frames 20-30) of each animal recording as this captures the background fluorescence of the neuron prior to stimulation. This adjusted ΔF/F_0_ value was corrected to be the percent change in fluorescence by the following equation: (ΔF/F_0_ - 1) * 100%. The average for each frame was calculated and plotted overtime with the standard error measure for each 0.1 second. For calcium imaging, the average value for each of these periods was plotted along with the SEM for each group. The maximum change in fluorescence values was compared to one another by One-way ANOVA with Tukey’s Multiple Comparisons test for parametric data and a Kruskal-Wallis H Test for non-parametric data (the pre-stimulus period is often non-parametric). Comparisons of imaging data within a sample (pre-stimulus to stimulus) was compared using a paired Student’s t-test for parametric data or a Wilcoxon signed rank test for non-parametric, paired data and Mann-Whitney test for non-parametric unpaired data. If multiple t-tests were used within a graph, Bonferroni’s correction is applied to adjust for multiple comparisons. At least ten animals were tested per condition, with one trial per animal.

### Avoidance Drop Assay

For experiments where the animals were infected, the animals were exposed to *S. aureus* grown on 35 mm TSA and kanamycin 10 µg/mL plates or *E. coli* grown on 35 mm NGM/TSA plates for 3 or 24 hours at 25 °C prior to testing. These infected animals were then transferred into 10 μL of M9 (22 mM KH_2_PO_4_, 42 mM Na_2_HPO_4_, 85 mM NaCl, 1 mM MgSO_4_) on an unseeded NGM (60 mm petri dish, 51 mM NaCl, 10 mM peptone, 51 mM agar, 25 mM KPO_4_ pH 6.0, 1 mM MgSO_4_, 1 mM CaCl_2_, 13 µM cholesterol) plate and allowed to crawl around to clean themselves, before this transfer process was repeated two more times, for a total of three cleaning cycles. From there, 10 animals were transferred to a new NGM plate. An avoidance drop assay was performed as previously described (27). To the tail of a forward moving animal, a small drop (∼5 nL) of solution was delivered via a hand pulled 10 μL glass capillary tube. Upon exposure, the solution was drawn up the body of the animal by capillary action where it is sensed by the amphid neurons. The animal is then categorized as having one of two behaviors; no avoidance (the animal continues moving forward) or avoidance (the animal reverses by two body bends within 4 seconds of exposure). Animals were first exposed to the solvent control (solvent without the stimulus, water) before the stimulus (1 M glycerol in water or 10 mm copper chloride in water) was applied. The avoidance index was calculated on a per plate basis as the number of animals that avoided drops over the total number of animals tested. One way ANOVA among stimulations with Tukey’s multiple comparisons was used. At least 10 plates were tested per condition, each containing at least 10 animals.

### PaIN image acquisition and analysis

*E. coli* and *S. aureus* plates were made the day prior to imaging. More than 50 L4 animals were added to the bacterial plates. The animals were transferred to a 25 °C incubator for 24 hours. After 24 hours, animals were placed in 100 mM sodium azide on top of a 4% agar pad slide. Animals were placed in close proximity before adding a cover slip and lightly sealing an edge with nail polish. Slides were imaged within 15 minutes of preparation. Animals were imaged in BioTek Lionheart FX automated microscope with the 20x objective. Ten to 20 animals were imaged per treatment. To score ASH PaIN, 3 or more breaks in the ASH dendrite was scored as a 1. The scored animals were summed and divided by the total number of animals, then multiplied by 100 to get a percentage of animals with PaIN. The percentage of each biological replicate was graphed with SEM. An unpaired two-tail *t* test was used for analysis. A *p-*value ≤ 0.05 was considered significantly different from the control.

### Statistics and Sample Size

Data analysis was performed in GraphPad Prism (version 10.4.1). All data sets were first checked for normal distribution and homogeneity of variance. For two sample comparisons, a paired or unpaired Student’s two-tailed *t* test was selected for parametric data. For non-parametric data, a Wilcoxon signed rank test was used for paired data and Mann-Whitney test for unpaired data. If multiple t-tests were used within a graph, Bonferroni’s correction was applied to adjust for multiple comparisons, by dividing the alpha value 0.05 by the number of comparisons. For three or more comparisons, a One-way ANOVA with Tukey’s Multiple Comparisons test for parametric data and a Kruskal-Wallis H Test for non-parametric data was used. Appropriate sample size was selected based on a power analysis of preliminary data. For calcium imaging, at least ten animals were tested per condition, with one trial per animal. For the avoidance assays, at least 10 plates were tested per condition, each containing at least 10 animals. For PaIN imaging, 2-3 biological replicates were captured with 10-20 animals per replicate.

## Results

### *S. aureus* infection impairs ASH activation by aversive stimuli

To define how ASH neuronal activity changes after *S. aureus* infection, we visualized calcium concentration in ASH *in vivo* using the genetically encoded calcium indicator GCaMP3 (26). We infected animals for 3 hours on agar plates of *S. aureus* (infected) or *E. coli* (noninfected) prior to ASH stimulation and recording (**Figure 1A**). Noninfected *E. coli* controls exhibited a typical increase in cytoplasmic calcium following a 10-second stimulation with glycerol, a well-known aversive stimulus that activates ASH (**Figure 1B and C, Supplemental Video 1, Supplemental Figure 1A**) (16, 26). In contrast, *S. aureus-*infected animals showed greatly impaired Ca^2+^ transients (**Figure 1B and C, Supplemental Video 2, Supplemental Figure 1B and C**). This result suggested that *S. aureus* infection disables the ASH response to glycerol.

**Figure 1:**
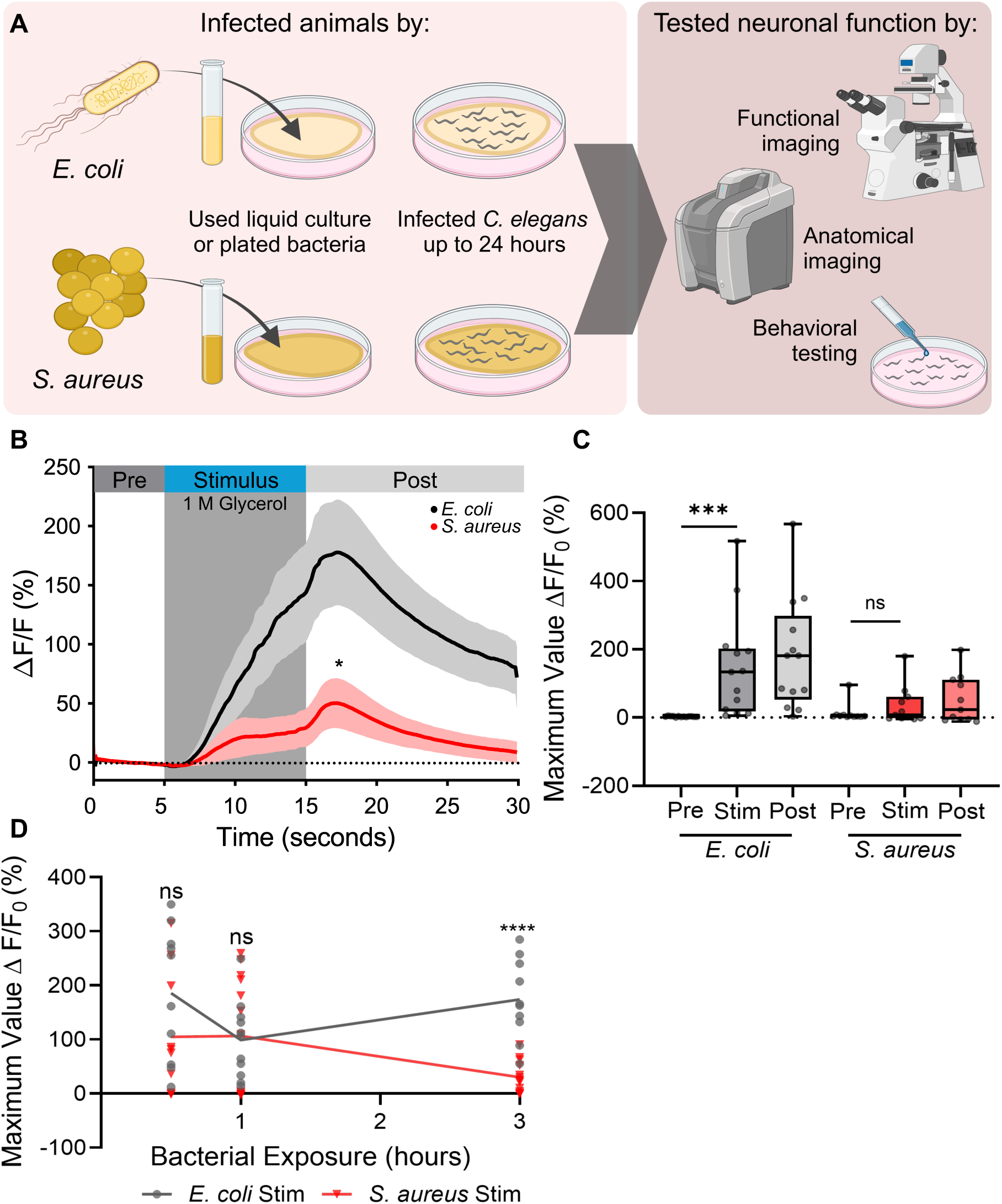
*S. aureus* infection impairs ASH neurons sensation and activity. **A)** *C. elegans* were infected with *E. coli* (OP50) or *S. aureus* (SH1000) for varying lengths of time before the function of ASH neurons was characterized by behavioral testing, anatomical neural imaging or functional neural imaging Created in BioRender. DiLoreto, E. (2025) https://BioRender.com/r16p000. **B)** Normalized change in GCaMP3 fluorescence over time. Animals were infected with *E. coli* or *S. aureus* for 3 hours. 10 sec exposure to 1 M glycerol (grey box). Data are average with SEM (shaded areas next to curves), *E. coli* n = 13, *S. aureus* n = 11. (*p = 0.0218 Maximum Value of Fluorescence during stimulation of 3 hour (hr) infected *E. coli* and *S. aureus* animals, Mann-Whitney t-test post to post). **C)** Maximum peak values of fluorescence changes during Pre (0-5 seconds (sec)), Stimulus (Stim) (5-15 sec), Post (15-30 sec) periods. Box and whisker plot with minimum and maximum values, median indicate. (*E. coli* ***p = 0.0002, *S. aureus* ^ns^*p* = 0.4648, Wilcoxon paired t-test). **D)** Percent change graphs of ASH imaging in infected animals at 0.5, 1, or 3 hours, stimulus of 1 M glycerol applied between 5-15 seconds. Each dot represents one individual animal. Line plots mean through each timepoint. (0.5 hr ^ns^*p* = 0.1454, 1hr ^ns^*p* = 0.8464, 3hr ****p < 0.0001, Unpaired t-test with Bonferroni’s correction, original *p ≤ 0.05, corrected *p ≤ 0.0167).

To determine the duration of infection necessary to disable the ASH response to glycerol, we performed infections for 0.5, 1, and 3 hours prior to ASH stimulation and calcium imaging. Animals that were infected for 0.5 and 1 hours were indistinguishable from noninfected controls, whereas animals infected for 3 hours showed an impaired response (**Figure 1D, Supplemental Figure 1D-F**). We concluded that impairment of the ASH response to glycerol requires longer than 1 hour but less than 3 hours of infection. Together, these data indicated that *S. aureus* infection disables the glycerol response of ASH over a long timeframe. This led us to directly investigate the effect of *S. aureus* on ASH neurons in the absence of noxious stimuli.

### ASH neurons depolarize during *S. aureus* exposure

To determine the effect of short *S. aureus* exposure on ASH activity, we visualized the calcium concentration in ASH neurons of animals exposed to *S. aureus* in its conditioned media. During the exponential (log) growth phase, the population experiences rapid growth until limiting media nutrients are depleted, at which time stationary phase is initiated (28). Because *S. aureus* physiology, including the expression of virulence factors, is influenced by population density (29), we compared exponential-phase to stationary-phase cultures of *S. aureus*. Prior to immobilization in the microfluidic device to view calcium activity, animals were reared under normal laboratory conditions. Once in the microfluidic device, they were exposed to a culture of log-phase or stationary-phase *S. aureus* grown under hypoxic conditions (**Figure 2A**). As control, a population of animals was exposed to Tryptic Soy Broth (TSB) without *S. aureus.* Animals exposed to TSB alone or log-phase *S. aureus* showed ASH activation, as indicated by a calcium transient (**Figure 2B and C**). In contrast, stationary-phase *S. aureus* did not activate ASH, but rather suppressed the calcium signal (**Figure 2B and C**). This result suggested that stationary phase *S. aureus* inhibits ASH through an unknown mechanism.

**Figure 2:**
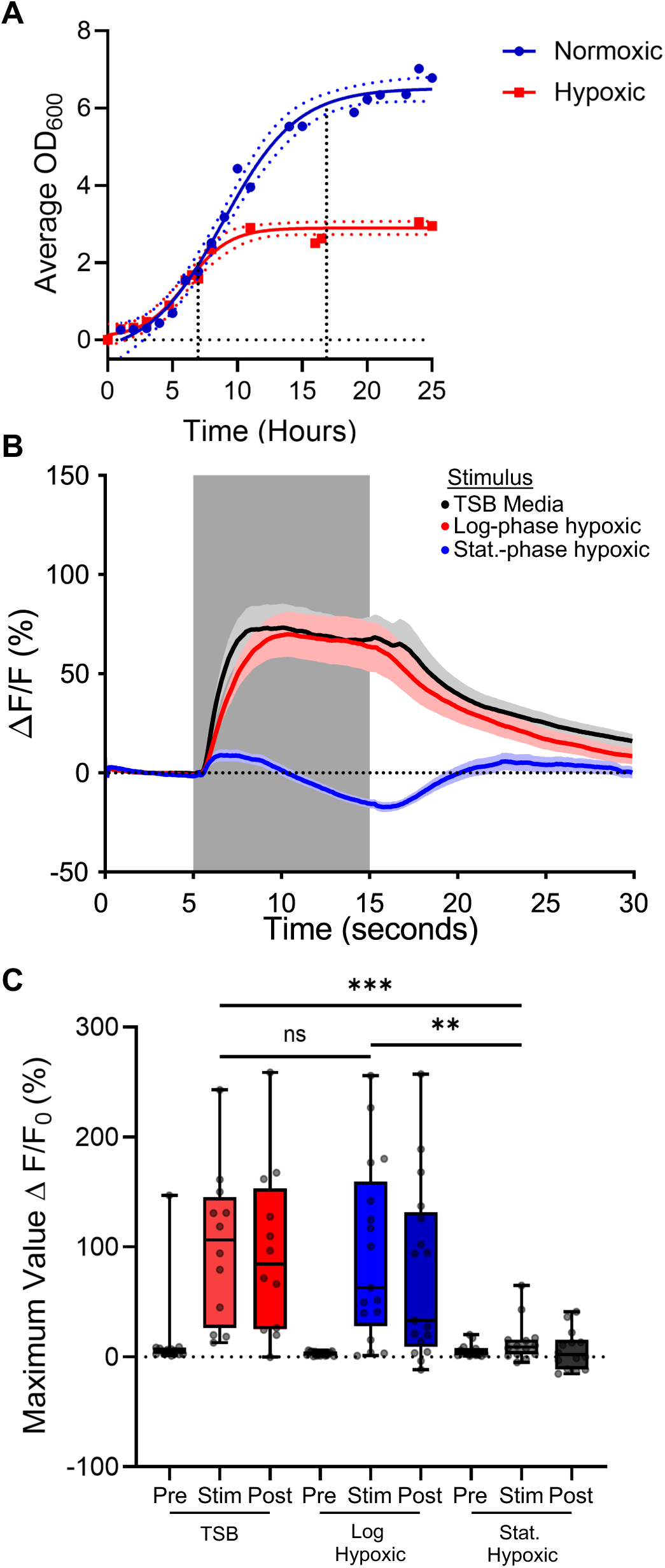
ASH neurons can sense changes in *S. aureus* culture. **A)** Normoxic (blue) and hypoxic (red) growth curve of *S. aureus* with 3 replicated per time point. Log phase first dashed line ∼6 hours, stationary phase second dashed line 16 – 18 hours. **B)** Response of ASH neuron to Log- (red) or stationary- (blue) phase hypoxic *S. aureus* compared with TSB stimulus (black) after application of a neutral S. Basal buffer. Trace is average response of > 10 animals, SEM indicated with shaded region. **C)** Maximum peak values of fluorescence changes during Pre (0-5 sec), Stim (5-15 sec), Post (15-30 sec). Plotted Min to Max values with median. (TSB Stim to Log Stim ^ns^p > 0.9999, TSB Stim to Stationary Stim ***p = 0.0009, Log Stim to Stationary Stim **p = 0.0028, Kruskal-Wallis with Dunn’s multiple comparisons).

### ASH inhibition depends on *S. aureus* growth conditions

*S. aureus* is a facultative anaerobe and has different metabolic, proteomic, and genetic profiles depending on the concentration of oxygen in culture (30–33). Under our previous hypoxic culture condition, the pH of the *S. aureus* media dropped from 7 to ∼4, consistent with the known fermentative metabolism of *S. aureus* (**Supplemental Figure 2A**) (32). To determine the effect of *S. aureus* culture oxygen concentration affects its ability to induce the ASH response, we cultured *S. aureus* in normal and low oxygen conditions prior application to the animals. We cultured *S. aureus* under normoxic and hypoxic conditions to both mid-log and stationary phases (**Figure 3A and D**). Normoxic culture did not cause the pH of the culture to decrease as from 7 as dramatically as the hypoxic growing conditions, allowing us to disentangle stationary phase from pH during ASH visualization (**Supplemental Figure 2A**). To exclude the effects of TSB media, the animals were first acclimated to unconditioned TSB media before the stimulus of *S. aureus*. We determined that with this “pre-activation” to TSB media, the ASH neurons were still capable of responding to a novel stimulus (**Supplemental Figure 2B and C**), showing that prior stimulation with TSB does not desensitize ASH to noxious stimuli. We extended the stimulus time to capture potentially late changes in neural activation.

**Figure 3:**
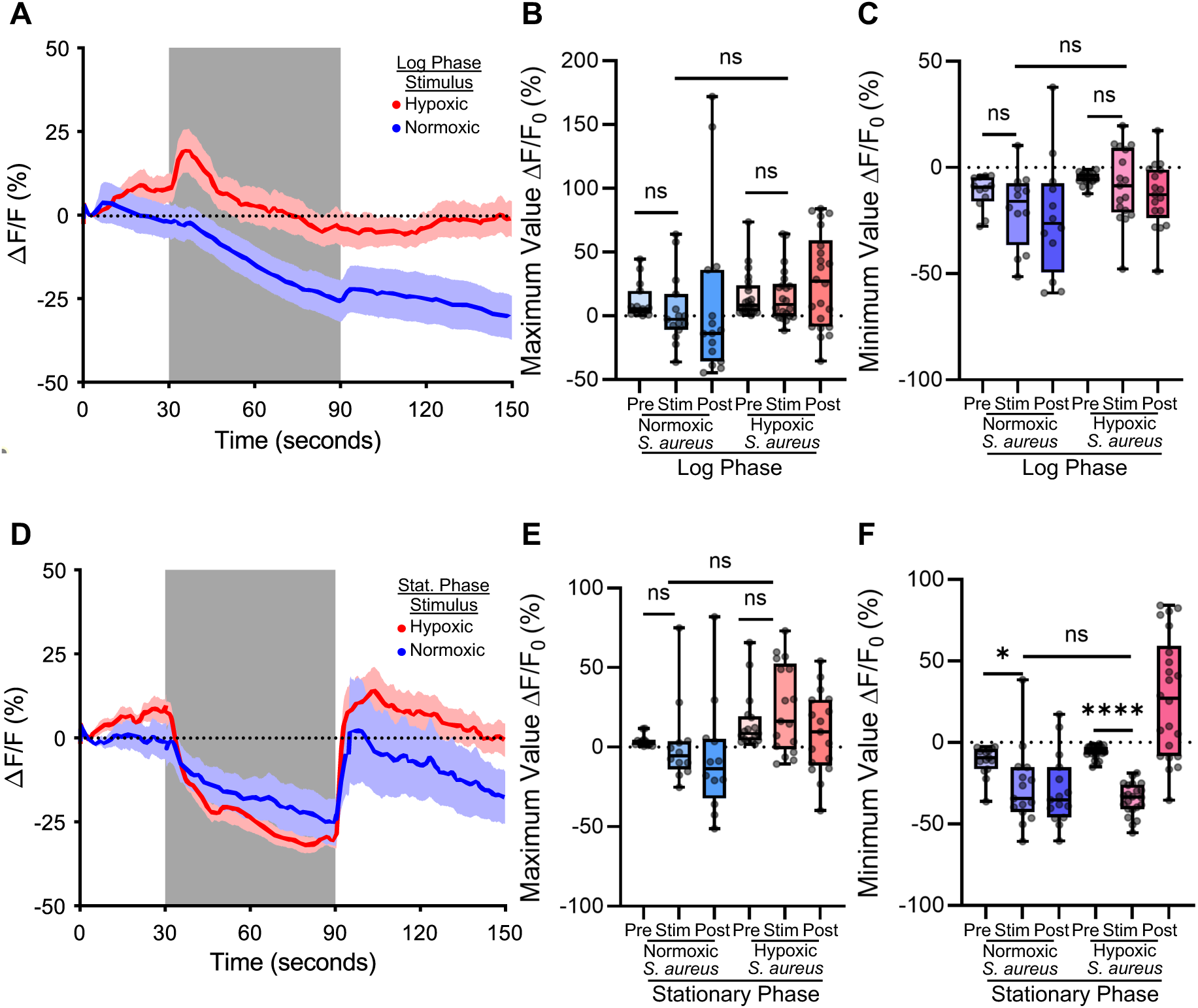
Growth-phase dependent inhibition of ASH activity by *S. aureus* activity. **A)** Log-phase normoxic (blue) and hypoxic (red) *S. aureus* exposed to ASH neuron (during grey bar) pre-exposed to TSB. Trace is average change in fluorescence intensity normalized to baseline fluorescence, shaded region is SEM. **B)** Maximum value of fluorescence intensity during Pre stimulus period (0-30 sec), Stimulus period (30-90 sec), and Post stimulus period (90-150 sec) of log phase *S. aureus* stimulation. (Normoxic Pre to Stim Wilcoxon ^ns^p = 0.3013; Hypoxic Pre to Stim Wilcoxon ^ns^p = 0.2247; Normoxic Stim to Hypoxic Stim Mann-Whitney ^ns^p = 0.0183). **C)** Minimum value of fluorescence intensity during Pre, Stimulus and Post periods of log phase *S. aureus* stimulation. (Normoxic Pre to Stim Paired t-test ^ns^p = 0.0337; Hypoxic Pre to Stim Paired t-test ^ns^p = 0.4164; Normoxic Stim to Hypoxic Stim Unpaired t-test ^ns^p = 0.1053). **D)** Stationary-phase normoxic and hypoxic *S. aureus* exposed to ASH neuron (during grey bar) pre-exposed to TSB. Trace is the average change in fluorescence intensity normalized to baseline fluorescence, shaded region is SEM. **E)** Maximum value of fluorescence intensity during Pre, Stimulus and Post periods of stationary phase *S. aureus* stimulation. (Normoxic pre to stim Wilcoxon p = 0.3880, Hypoxic pre to stim Wilcoxon p = 0.0730, Normoxic stim to Hypoxic stim Mann-Whitney p = 0.0386). **F)** Minimum value of fluorescence intensity during Pre, Stimulus and Post periods of stationary phase *S. aureus* stimulation. (Normoxic pre to stim Wilcoxon *p = 0.0103, Hypoxic pre to stim Wilcoxon ****p < 0.0001, Normoxic stim to Hypoxic stim Mann-Whitney ^ns^p = 0.5306). Box and whiskers plot minimum to maximum values with median indicated. Each individual point is the value of one animal. (All statistics with Bonferroni’s multiple comparison correction, original *p ≤ 0.05, corrected *p ≤ 0.0167).

In animals exposed to normoxic mid-log phase *S. aureus,* calcium levels trended lower than controls over 60 seconds of exposure and remained low after *S. aureus* removal (**Figure 3A**). However, this trend did not reach statistical significance either by comparing the maximal fluorescence change values for each animal during different phases of stimulation or by comparing the minima (**Figure 3B and C**). In contrast, in animals exposed to hypoxic mid-log *S. aureus* calcium levels to initially trended higher followed by a downward trend that slowly recovered after *S. aureus* removal; however, these trends also did not reach statistical significance (**Figure 3B and C**). We concluded that mid-log *S. aureus* caused slight decrease in cytosolic calcium in ASH neurons.

Stationary phase *S. aureus* suppressed calcium concentration with faster kinetics than exponential phase (**Figure 3D – F**). [Say something about statistical significance?] Normoxic and hypoxic cultures exhibited similar kinetics and magnitudes of ASH suppression (**Figure 3D – F**). Thus, in exponential phase, normoxic *S. aureus* was slightly more effective at reducing ASH activity than hypoxic *S. aureus,* while in stationary phase both conditions were equally effective. These data suggest that stationary phase increases the ability of *S. aureus* cultures to inhibit ASH, regardless of culture oxygen and media pH.

### ASH inhibition is associated with pathogen-induced neurodegeneration (PaIN)

The two ASH cell bodies in *C. elegans* reside near the nerve ring, in the posterior end of the head, and project one dendrite each to openings near the mouth (**Figure 4A**) (16). In our previous calcium recordings, which focused on the neuron cell body, we noticed that infected animals frequently showed defective dendrites. The observed phenotype, which we termed pathogen-induced neurodegeneration (PaIN), evidenced dendrite gaps and beading as revealed by GFP-labelled ASH *in vivo* (**Figure 4A lower**). After 3 hours of *S. aureus* infection, we observed a slight and statistically nonsignificant increase in this phenotype compared to noninfected controls (**Figure 4B**). The PaIN phenotype worsened considerably by 24 hours of infection to significantly different levels than noninfected controls (**Figure 4C**). Control experiments showed that 24 hours of starvation or OP50 feeding on TSA media used in the *S. aureus* infection assays do not induce PaIN, ruling out the possibility that *S. aureus-*induced intestinal destruction (i.e. lack of nutrition) or the culture medium induce PaIN by themselves (**Supplemental Figure 3**). Thus, ASH PaIN is specifically caused by *S. aureus* and takes place between 3 and 24 hours.

**Figure 4:**
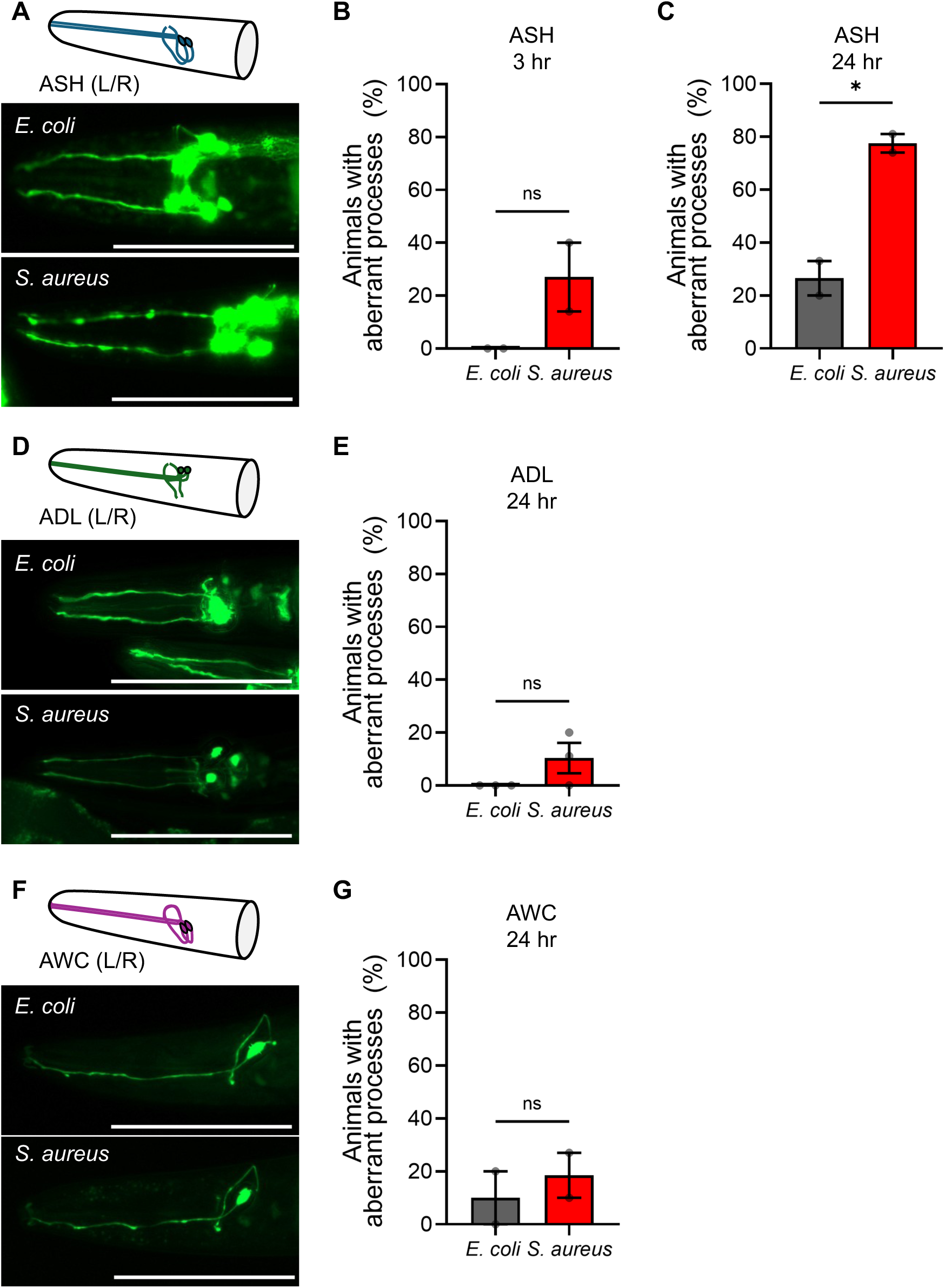
Pathogen Induced Neurodegeneration (PaIN) disproportionately affects ASH Sensory Neurons. **A)** Cartoon of ASH neurons alongside representative epifluorescence micrographs of ASH::GCaMP (CX10979) animals fed *E. coli* (top) or infected with *S. aureus* (bottom) for 24 hours at 25 °C. Scale bar = 100 μm. **B)** Quantification of ASH PaIN in animals infected for 3 hrs (^ns^p = 0.1734). **C)** Quantification of ASH PaIN in animals infected for 24 hrs (*p = 0.0203). **D)** Representative cartoon of ADL neurons alongside representative epifluorescence. ADL::GCaMP (OH14884) animals fed *E. coli* (top) or infected with *S. aureus* (bottom) for 24 hrs. Scale bar = 100 μm. **E)** Quantification of ADL PaIN in animals infected for 24 hrs (^ns^p = 0.1475) **F)** Cartoon of AWC neurons alongside representative epifluorescence micrographs of AWC::GCaMP (CX10536) animals head fed *E. coli* (top) or infected with *S. aureus* (bottom) for 24 hrs. Scale bar = 100 μm. **G)** Quantification of AWC PaIN in animals infected for 24 hrs (^ns^p = 0.5836). Average ± SEM of 2-3 biological replicates each with n = 10-25 animals. (Unpaired two-tailed t-test).

ASH is one of 12 amphid-ciliated sensory neurons located in the *C. elegans* that are exposed to the environment (34). To determine the specificity of the *S. aureus-*caused PaIN phenotype to ASH neurons, we examined neighboring amphid-ciliated sensory neurons. We examined ADL (Amphid Dual Ciliated Ending L, left/right pair) sensory neurons, which have similar position and project dendrites to openings next to the mouth, similar to ASH (**Figures 4D**) (15). After 24 hours of infection, we did not observe an increase in the number of defective ADL dendrites (**Figure 4E**). Similarly, we examined AWC (Amphid Wing Neuron C left/right pair) sensory neurons, which show similar position to ASH and ADL, but display wing-like cilia and that do not reach the exterior by the mouth (**Figure 4F**) (34). Like ADL neurons and unlike ASH neurons, we did not observe an increase in defective AWC dendrites after 24 hour infection with *S. aureus* (**Figure 4G**). We concluded that *S. aureus* induces the degeneration of ASH but not ADL and AWC neurons, suggesting that *S. aureus* induces PaIN specifically in ASH neurons.

### PaIN is *S. aureus*-specific

Several bacterial species can be found in natural association with *C. elegans.* Efforts to define a minimal set as a natural intestinal microbiota for *C. elegans* have yielded a reference collection of 12 species, named the CeMbio (*C. elegans* Microbiome) (35). In previous work, we showed that these isolates overall do not exhibit virulence against immunocompetent *C. elegans.* Instead, a subset shows virulence against specific immunocompromised mutants, a phenomenon termed cryptic virulence (36).To evaluate the ability of the CeMbio bacteria to cause ASH PaIN, we mono-associated *C. elegans* with each isolate. While *E. coli* controls showed no PaIN and *S. aureus-*infected animals showed significantly increased PaIN, animals in mono-association with most of the CeMbio bacteria did not show PaIN (**Figure 4C and 5**). However, animals associated with *Pseudomonas lurida* MYb11 showed slight, but significant, induction of PaIN after 24 hours (**Figure 5**). This result suggests that *S. aureus*-induced PaIN may be an extreme manifestation of a phenotype that is induced by distinct bacteria that naturally associate with *C. elegans*.

**Figure 5:**
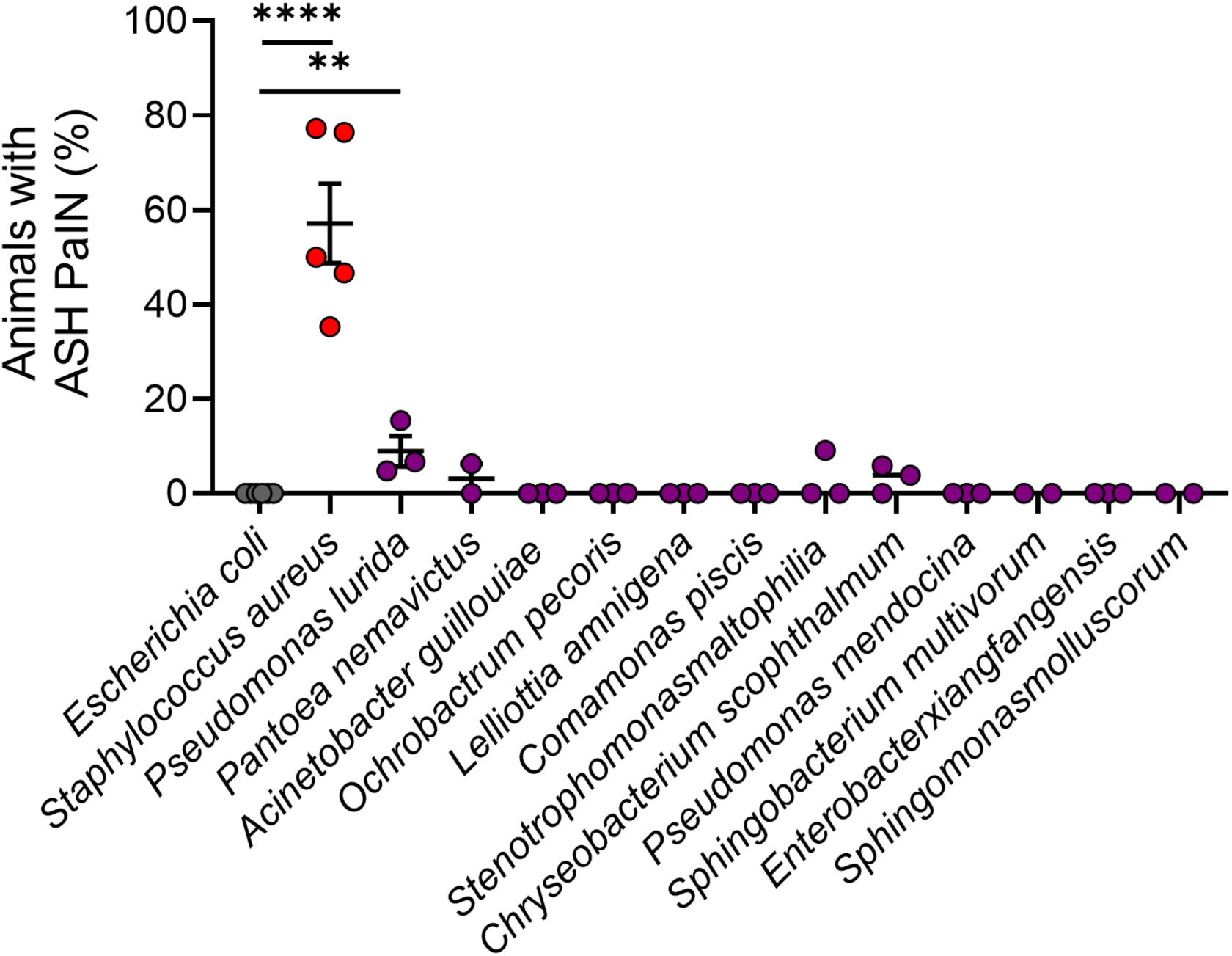
*C. elegans* microbiota members do not induce ASH PaIN, except for *P. lurida*. Quantitative percentage of animals showing ASH PaIN. Animals fed *E. coli*, infected with *S. aureus*, or associated with individual CeMbio members for 24 hrs. Average ± SEM, 3-5 biological replicates, n = 10 - 25 animals per biological replicate. (*E. coli* to *S.* aureus ****p < 0.0001, *E. coli* to *P. lurida \*\**p = 0.0042, Unpaired 2-tail t-test with Bonferroni correction, original **p ≤ 0.01, ****p ≤ 0.0001, corrected **p ≤ 0.005, ****p ≤ 0.00005).

### PaIN partially requires HLH-30/TFEB

Neurodegeneration in *C. elegans* and mammals may occur as a result of programmed cell death. In addition to apoptosis, neuronal death has been described to occur in *C. elegans* through necrosis and ferroptosis (37–39). To define their roles in *S. aureus*-induced ASH PaIN, we examined PaIN in mutants that are defective in each cell death modality. *ced-3* mutants, defective in apoptosis, did not show altered PaIN (**Figure 6A**) (37, 40, 41). Neither did *asp-4* or *itr-1* mutants (defective in necrosis) (**Figure 6B**) (42–45) nor *fat-3* mutants (defective in ferroptosis) (**Figure 6C**) (46–49). Thus, ASH PaIN is unlikely to proceed through apoptosis, necrosis, or ferroptosis.

**Figure 6:**
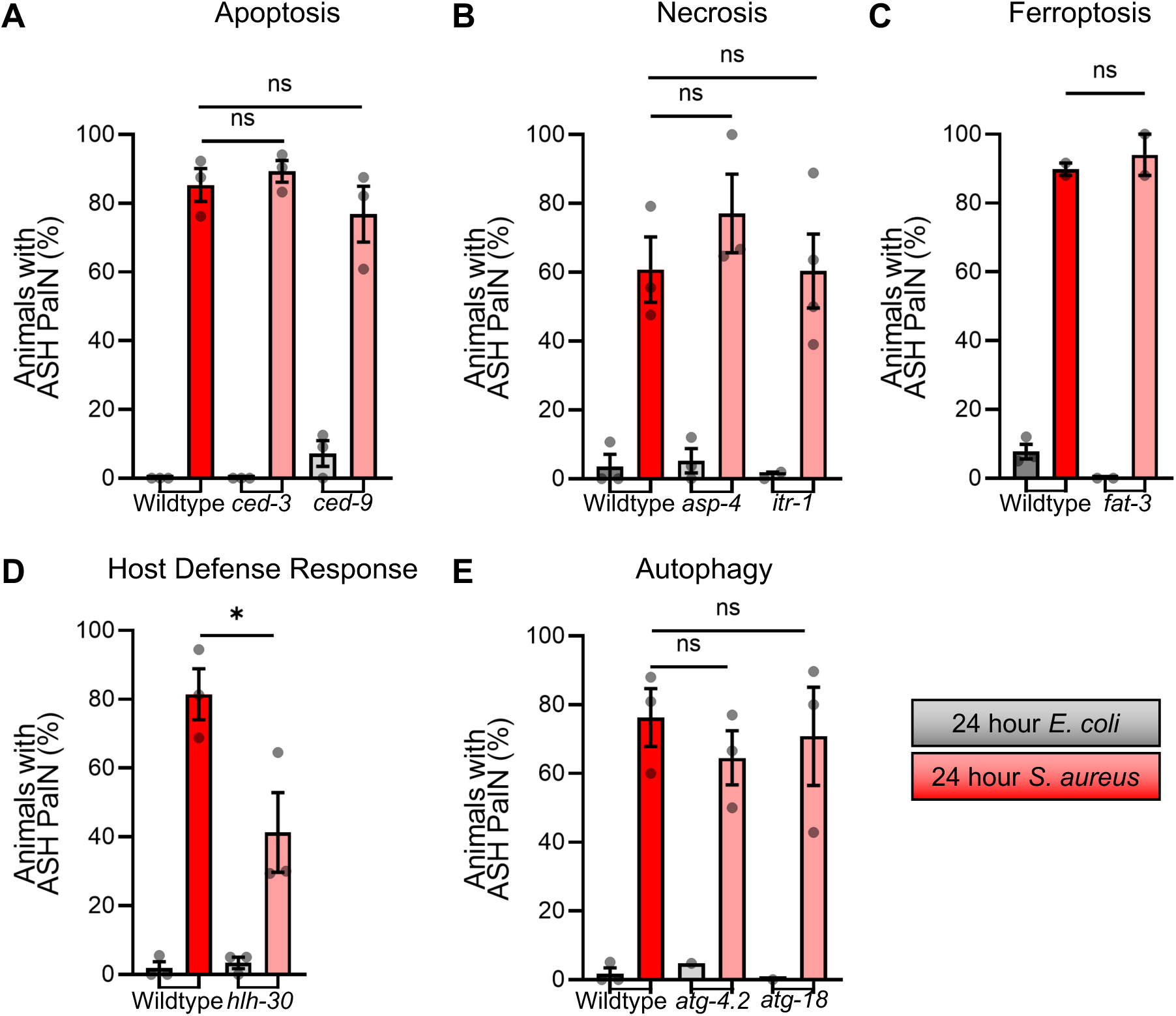
Necrosis, apoptosis, ferroptosis, and autophagy genes are dispensable for ASH PaIN. **A)** ASH PaIN tested in relation to apoptosis mechanisms by *ced-3* and *ced-9* loss of function mutants (*ced-3* ^ns^p = 0.5260, *ced-9* ^ns^p = 0.4187). **B)** ASH PaIN tested in relation to necrosis mechanisms by *asp-4* and *itr-1* loss of function mutants (*asp-4* ^ns^p = 0.7517, *itr-1* ^ns^p = 0.9798). **C)** ASH PaIN tested in relation to ferroptosis mechanisms by *fat-3* loss of function mutant (*fat-3* ^ns^p = 0.5756). **D)** ASH PaIN tested in relation to autophagy/lysosomal biogenesis mechanisms by *hlh-30* loss of function mutant (*hlh-30* ^ns^p = 0.0434). **E)** Autophagy pathways investigated in relation to ASH PaIN by *atg-4.2* and *atg-18* mutants (*atg-4.2* ^ns^p = 0.3663, *atg-18* ^ns^p = 0.7581). Average ± SEM from two – three biological replicates; n = 15 - 25 animals per biological replicate. (****p ≤ 0.0001, ***p ≤ 0.001, **p ≤ 0.01, *p ≤ 0.05. Unpaired two-tail t-test *p ≤ 0.05, Bonferroni’s correction applied to graphs with two statistical comparisons, corrected p ≤ 0.025).

In mammals, neurons can also die by autophagy- and lysosome-mechanisms (50, 51). To determine how disruption of autophagy and lysosomal biogenesis may affect ASH PaIN, we used *hlh-30/TFEB* mutants. Loss of *hlh-30* decreased the frequency of PaIN by about 50% compared to wildtype (**Figure 6D**), suggesting that HLH-30/TFEB is partially required for full ASH PaIN.

HLH-30 and TFEB are well-known as positive regulators of autophagy gene expression (10). Moreover, autophagy genes and autophagy itself are induced by HLH-30/TFEB during *S. aureus* infection, and the loss of autophagy enhances *C. elegans* susceptibility to *S. aureus-*mediated killing (52). To test if autophagy plays a role in ASH PaIN, we examined PaIN in loss-of-function mutants *atg-4.2* and *atg-18,* two autophagy genes that function in neurons (53, 54). However, both *atg-4.2* and *atg-18* mutants were indistinguishable from wildtype (**Figure 6E**), suggesting that autophagy is dispensable for ASH PaIN. Mutants that disrupt lysosome-mediated cell death are not available, precluding our investigation of that mechanism. Altogether, these data suggest that HLH-30/TFEB promotes ASH PaIN through an autophagy-independent mechanism that is also independent of major cell death pathways.

### *S. aureus* infection disrupts behavior

Copper (Cu^2+^) and glycerol are sensed by ASH neurons, which feed into the motor program to promote evasion of these aversive stimuli (16). To define the physiological relevance of the inhibition of ASH function and the induction of PaIN by *S. aureus,* we measured Cu^2+^ aversion by infected animals. Cu^2+^ is detected by both ADL, which can sense heavy metals but do not exhibit PaIN, and ASH sensory neurons, which exhibit PaIN (**Figure 4C and E**) (55). Animals infected with *S. aureus* for 3 hours showed aversion to Cu^2+^ that was indistinguishable from noninfected controls (**Figure 7A**). In contrast, after 24 hours of infection, infected animals showed a marked reduction in Cu^2+^ avoidance compared with noninfected controls (**Figure 7B**). This result suggested that aversive behaviors are negatively impacted by *S. aureus* infection over a timescale that correlates with ASH PaIN severity.

**Figure 7:**
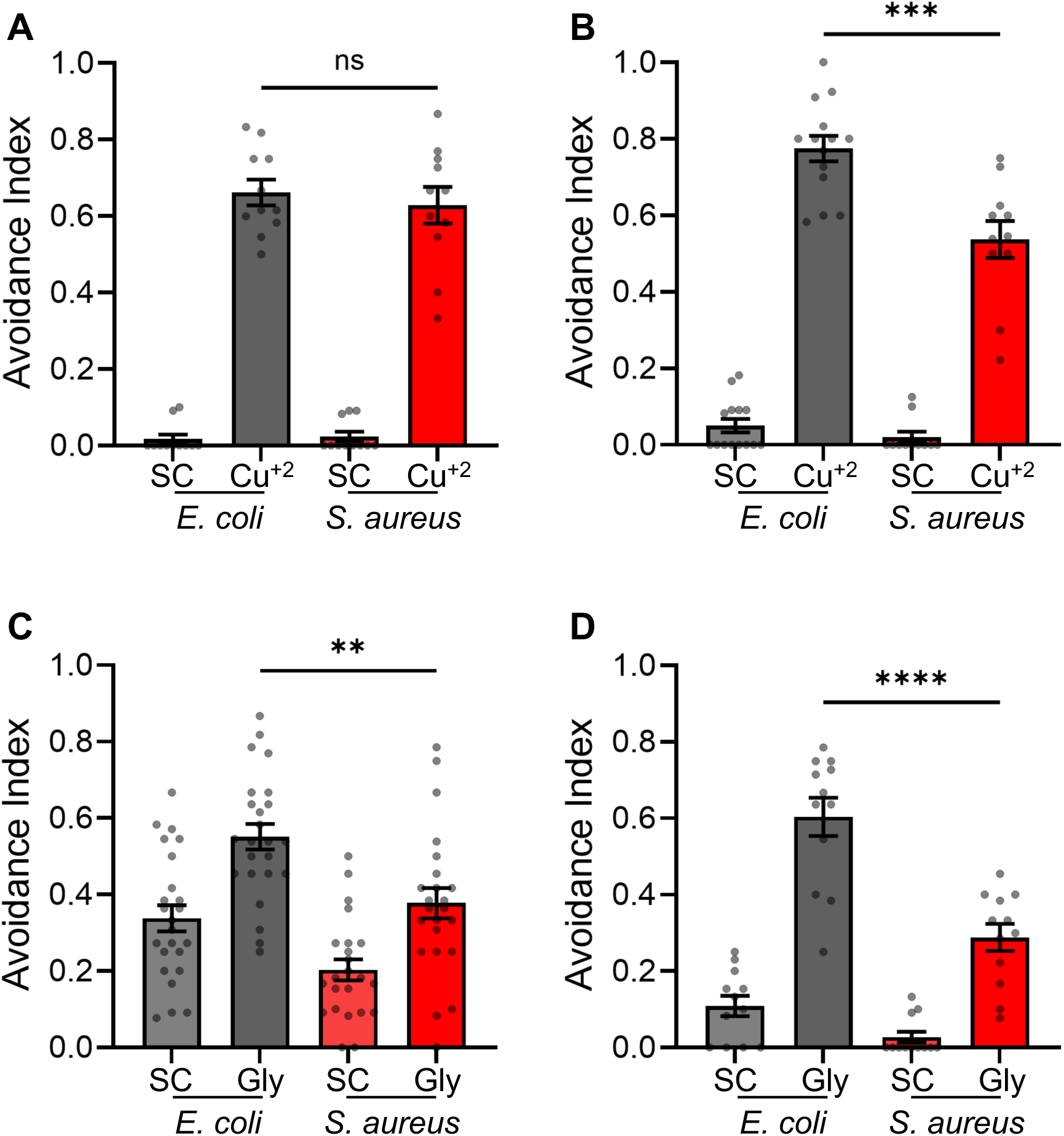
*S. aureus* infection impacts ASH associated behaviors. **A,B)** Drop Avoidance Assay of animals fed *E. coli* grown on TSA plates compared to *S. aureus* grown on TSA plates with 10 µg/mL kanamycin for **A)** 3 or **B)** 24 hours at 25°C before exposure to a drop of water (solvent control, SC) or 10 mM copper chloride (Cu^+2^). (3 hrs ^ns^p = 0.5727, 24 hrs ***p = 0.0004). **C, D)** Drop Avoidance Assay of animals fed *E. coli* grown on TSA plates compared to *S. aureus* grown on TSA plates with 10 µg/mL kanamycin for **C)** 3 or **D)** 24 hours at 25°C before exposure to a drop of water (solvent control, SC) or 1 M glycerol (Gly). (3 hrs **p = 0.0017, 24 hrs ****p < 0.0001). **A, B, C, D)** Average ± SEM. Each point is one plate of animals each with n > 10 animals per plate. Students two-tailed t-test.

To further test this conclusion, we used glycerol avoidance as a second ASH-dependent aversive behavior. In this assay, defective avoidance was apparent by 3 hours of infection (**Figure 7C**) and worsened by 24 hours (**Figure 7D**). These data are consistent with the hypothesis that ASH PaIN impairs ASH function and behavior after *S. aureus* infection.

## Discussion

The ASH neurons, which are amphid sensory neurons directly connected to the animal’s external environment, have long been characterized as polymodal nociceptive neurons that respond to various aversive stimuli (16, 17, 27, 56). We found that *S. aureus* quickly represses the activity of *C. elegans* ASH neurons. This ASH repression was dependent on the late, stationary phase of *S. aureus* growth, suggesting that it may require a specific pathogen phenotype. As in mammals, sensory neurons emerge as mechanisms for pathogen detection in *C. elegans*, responding to metabolites and odors related to microbes. For instance, *Escherichia coli* produces propyl acetate and butyl acetate metabolites, which elicit a response in AWA (Amphid Wing Neuron A, left/right pair) to induce attraction (57). Additionally, *Pseudomonas aeruginosa* creates 1-undecene which triggers amphid neuron AWB (Amphid Wing Neuron B, left/right pair) activation, which causes attraction behaviors (58). More recently *Enterococcus faecalis* was shown to induce host defense gene expression through unidentified volatile compounds that function through AWA and perhaps AFD (Amphid Finger-like Endings D, left/right pair) (6). Further work will need to investigate which *S. aureus* metabolites causes the change in ASH activity.

Towards this objective, we challenged *S. aureus* growth conditions. Since oxygen availability has a major impact on metabolism in *S. aureus* (29) and both high- and low-oxygen cultures repressed ASH activity, such repression is unlikely to be the result of metabolic changes between culturing conditions. Similarly, since normoxic cultures did not modify the pH of the media, low pH does not appear to be the cause of ASH repression. Previous research has shown that ASH becomes activated, not inhibited, by low pH stimuli (17, 18). Further work is needed to define if soluble mediators produced by *S. aureus* inhibit ASH activity, and what their identity might be. Where we found a difference in ASH suppression was between exponential and stationary phase cultures of *S. aureus*. Previous studies have found that the pathogenicity of *S. aureus* may change throughout its growth (59). During late-exponential growth, there are more toxins produced and more toxin forming genes upregulated in *S. aureus* culture, and environmental factors can influence this regulation (59, 60). Several virulence factors are known to be induced in stationary phase in *S. aureus,* notably pore-forming toxins (61). Pore-forming toxins created by *S. aureus* have been shown to activate mammalian nociceptor neurons, raising the possibility that such virulence factors may differentially affect nociceptor neurons in nematodes and mammals (62). The roles of pore-forming toxins and other virulence factors in ASH repression will be a major focus for future work.

We also discovered that *S. aureus* infection causes neuron-specific degeneration. Infection by *S. aureus* results in structural changes specifically targeting the ASH neurons, which we term pathogen-induced neurodegeneration (PaIN). This PaIN manifests as breaks along the ASH dendrites, resembling the beaded dendrites (63) or blebbing (64) observed in other neurodegenerative conditions, even in mammals. ASH PaIN is detectable early, by 3 hours of infection, and becomes severe by 24 hours, at which time ASH-dependent aversive behavior is defective. We have previously found that this time point was selected as preliminary, as prior investigations revealed increased ACh levels in *S. aureus* infected animals compared to *E. coli*-fed animals and consequently, the host defense response by this time (8). The PaIN phenotype we observe in ASH resembles other *C. elegans* neurodegeneration phenotypes, including degeneration of neurons including ASI (amphid single cilium neuron I) during *P. aeruginosa* infection (63), PVQ (posterior ventral process neuron Q) during mitochondrial loss (65), and PVD (posterior ventral process neuron D) during hyperactivation of NLP-29 peptides by an overactive immune response to fungal infection (64). *P. aeruginosa* infection can also cause PaIN in ASE (amphid single cilium neuron E) and AWC amphid neurons, which neighbor ASH (63, 66, 67). In contrast, we did not observe PaIN in AWC and other amphid neurons aside from ASH neurons during *S. aureus* infection. This suggests that ASH PaIN is not part of a general neurodegeneration during infection, but rather a specific interaction between *S. aureus* and ASH neurons. We have demonstrated that *S. aureus* causes morphological changes consistent with degeneration in ASH (but not two other sensory neurons that were tested) and weakens ASH-dependent aversive behavior, suggesting that the ASH neurons are functionally as well as morphologically impaired by infection. Further work will investigate how this disrupted ASH function impairs downstream neural communication since ASH are highly connected sensory neurons (68).

Genetic analysis suggests that *S. aureus-*induced ASH PaIN occurs independently of common cell death pathways, in contrast to certain mammalian neurodegenerative conditions. However, deletion of the host defense transcription factor HLH-30/TFEB reduced the penetrance of PaIN through an unknown mechanism that is unlikely to involve autophagy. Future investigations will uncover these mechanisms. HLH-30/TFEB is known as a master regulator of autophagy and lysosomal biogenesis (10, 69), and there is long-standing evidence of the involvement of autophagy-dependent and lysosome-dependent mechanisms in neurodegeneration in humans and other animal models (70–73). There is also a longstanding theory that microbial infection may precede neurodegeneration (74). Supporting this theory of infection-promoted neurodegeneration by host defense mechanisms, previous research shows that PVD dendrites are enriched with autophagosomes and fragmented microtubules during fungal infection and aging (63). However, this response in PVD neurons were triggered by ligand binding to the G protein-coupled receptor NPR-12, which ASH does not express (75). This might suggest that the impact of HLH-30/TFEB occurs outside of ASH in some of the downstream neurons in the neuro-immune circuit. Further exploration of the start of this host-immune pathway will be important as TFEB has been implicated in human neurodegenerative diseases and is a therapeutic target for multiple neurodegenerative diseases (76, 77). Involved in this ASH neurodegeneration mechanism is the HLH-30/TFEB pathway, which impacts the degree of ASH PaIN. Future work will explore if ASH PaIN could be prevented or treated after *S. aureus* infection. Future studies will investigate whether ASH-mediated ACh release is the signal that activates the muscarinic-WNT host defense pathway. This information provides evidence that supports the specificity and important regulation of the neuro-immune response along the gut-brain axis.

## Conclusion

The biological logic of ASH PaIN during infection by *S. aureus* remains unclear. Our results showing that *Pseudomonas lurida* causes mild ASH PaIN, suggesting that the morphological and functional impairment of ASH may be relevant in the *C. elegans* native environment. We have previously found that both *P. lurida* and *S. aureus* decrease survival among *C. elegans* (36). In one scenario (bacteria-first model), bacteria (pathogens) may have evolved the ability to disrupt ASH neurons, thus promoting favorable host behaviors. A lack of aversion towards pathogen-derived molecules could favor the ingestion of the pathogen, promoting infection and dispersion. In a likelier scenario (host-first model), the host may have evolved the ability to suppress evasive behaviors by eliminating neurons that trigger them. This may be advantageous under conditions where the host cannot escape pathogens by evading chemical signals, such as in *S. aureus* infection plates. Under these conditions, it may benefit the host to become insensitive to some environmental cues and charge on in a linear direction with the chance to leave a noxious environment. This may explain why *S. aureus-*infected ASH-animals forged up the plate sides even as they died from desiccation. Further research is needed to test the bacteria-first and host-first models and their implications for higher organisms.

## List of Abbreviations

*C. elegans*: *Caenorhabditis elegans*
*S. aureus*: *Staphylococcus aureus*
*E. coli*: *Escherichia coli*
*P. lurida*: *Pseudomonas lurida*
ASH: Amphid Single Ciliated Neuron H
ADL: Amphid Dual Ciliated Neuron L
AWC: Amphid Wing Ciliated Neuron L
ACh: acetylcholine
CeMbio: C. elegans Microbiome bacteria
Cu^2+^: copper
HLH-30/TFEB: helix-loop-helix protein 30 / transcription factor EB

## Declarations

- Ethics approval and consent to participate- not applicable
- Consent for publication- not applicable
- Availability of data and materials- All data generated or analyzed during this study are included in this published article and its supplementary information files. All bacterial and nematode strains can be provided upon request.
- Competing interests- the authors declare no competing interests
- Funding- grants R01DC016058 (JS), R35GM149284 (JEI), R01GM101056 (JEI), T32AI095213 (XG) and R25GM113686 (XG) from the National Institutes of Health of the USA, and the Dr. Marcellette G. Williams Memorial Fund (JEI).

### Authors’ contributions

- EMD- conceptualization, investigation of calcium imaging of ASH and behavior, formal analysis, visualization, validation, writing-original draft, writing-review and editing.
- XG- investigation of PaIN phenotype in ASH, ADL, CEMbio strains and mutant mechanisms of PaIN, formal analysis, visualization, writing-original draft.
- KAW- investigation of PaIN phenotype in ASH and AWC, formal analysis, visualization.
- JSh- investigation of mutants involved in PaIN mechansism, formal analysis, visualization.
- JEI- conceptualization, formal analysis, visualization, funding acquisition, supervision, writing-review and editing.
- JSr- conceptualization, funding acquisition, supervision, writing-review and editing.
- All authors read and approved the final manuscript

## Acknowledgements

Some strains were purchased from the *Caenorhabditis* Genetics Center which is funded by NIH Office of Research Infrastructure Programs (P40 OD010440).

- Authors’ information (optional)- not applicable

## Supporting Information

**Supplemental Table 1:** *C. elegans* Strains Used.

| Strain Name | Genotype | Procured From | Figure in this Paper |
| --- | --- | --- | --- |
| N2 | Bristol Wildtype isolate | CGC | 7 |
| CX10536 | *kyEx2995[*str-2p::GCaMP2.2b*, *unc-122::GFP*]* | CGC | 4F, G |
| CX10979 | *N2;kyEx2865[*sra-6p::GCaMP3*]* | Bargmann Lab | 1B-D, 2, 3, Sup 1, Sup 2B-C |
| JIN2164 | *kyEx2865[*sra-6p::GCaMP3*]; *asp-4(ok2693)X** | This work | 6B |
| JIN2166 | *CX10979;MT4770 kyEx2865[*sra-6p::GCaMP3*]; *ced-9(n1950)I** | Irazoqui Lab | 6A |
| JIN2167 | *kyEx2865[*sra-6p::GCaMP3*]; *hlh-30(tm1978)IV** | This work | 6D |
| JIN2191 | *oyls14[*sra-6p::GFP + lin-15(+)*]V.; otls38 [*unc-47(del)p::GFP*]X* | CGC | 4A-C, 5, Sup 3 |
| JIN2192 | *oyls14[*sra-6p::GFP + lin-15(+)*]V; otls38[*unc-47(del)::GFP*]X* | Irazoqui Lab | 6 |
| JIN2231 | *itr-1(sa73)IV; oyIs14[*sra-6p::GFP + lin-15(+)*]V; otls38[*unc-47(del)::GFP*]X* | This work | 6C |
| JIN2233 | *ced-3(n717)IV; oyIs14[*sra-6p::GFP + lin-15(+)*]V; otls38[*unc-47(del)::GFP*]X* | This work | 6A |
| JIN2235 | *fat-3(wa22)IV; oyIs14[*sra-6p::GFP + lin-15(+)*]V; otls38[*unc-47(del)::GFP*]X* | This work | 6C |
| JIN2276 | *atg-4.2(ola316)IV; oyIs14[*sra-6p::GFP + lin-15(+)*]V; otls38[*unc-47(del)::GFP*]X* | This work | 6E |
| JIN2289 | *atg-18(gk378)V ; oyIs14[*sra-6p::GFP + lin-15(+)*]V; otls38[*unc-47(del)::GFP*]X* | This work | 6E |
| N2 | Bristol Wildtype isolate | CGC | 7 |
| OH14884 | *pha-1(e2123) III; him-5(e1490) V; otls646* | CGC | 4D, E |

**Supplemental Video 1: *C. elegans* ASH neuron after 3 hours of *E. coli* feeding**

*C. elegans* labeled with GCaMP3.0 in ASH neurons exposed to OP50 *E. coli* on TSA plates for 3 hours. Animals were exposed to aversive chemical 1 M glycerol starting at second 5 for 10 seconds.

**Supplemental Video 2: Activity in *C. elegans* ASH neuron decreased after 3 hours of *S. aureus* infection**

*C. elegans* labeled with GCaMP3.0 in ASH neurons exposed to SH1000 *S. aureus* on TSA + 10 µg/mL kanamycin plates for 3 hours. Animals were exposed to aversive chemical 1 M glycerol starting at second 5 for 10 seconds.

**Supplemental Figure 1:**
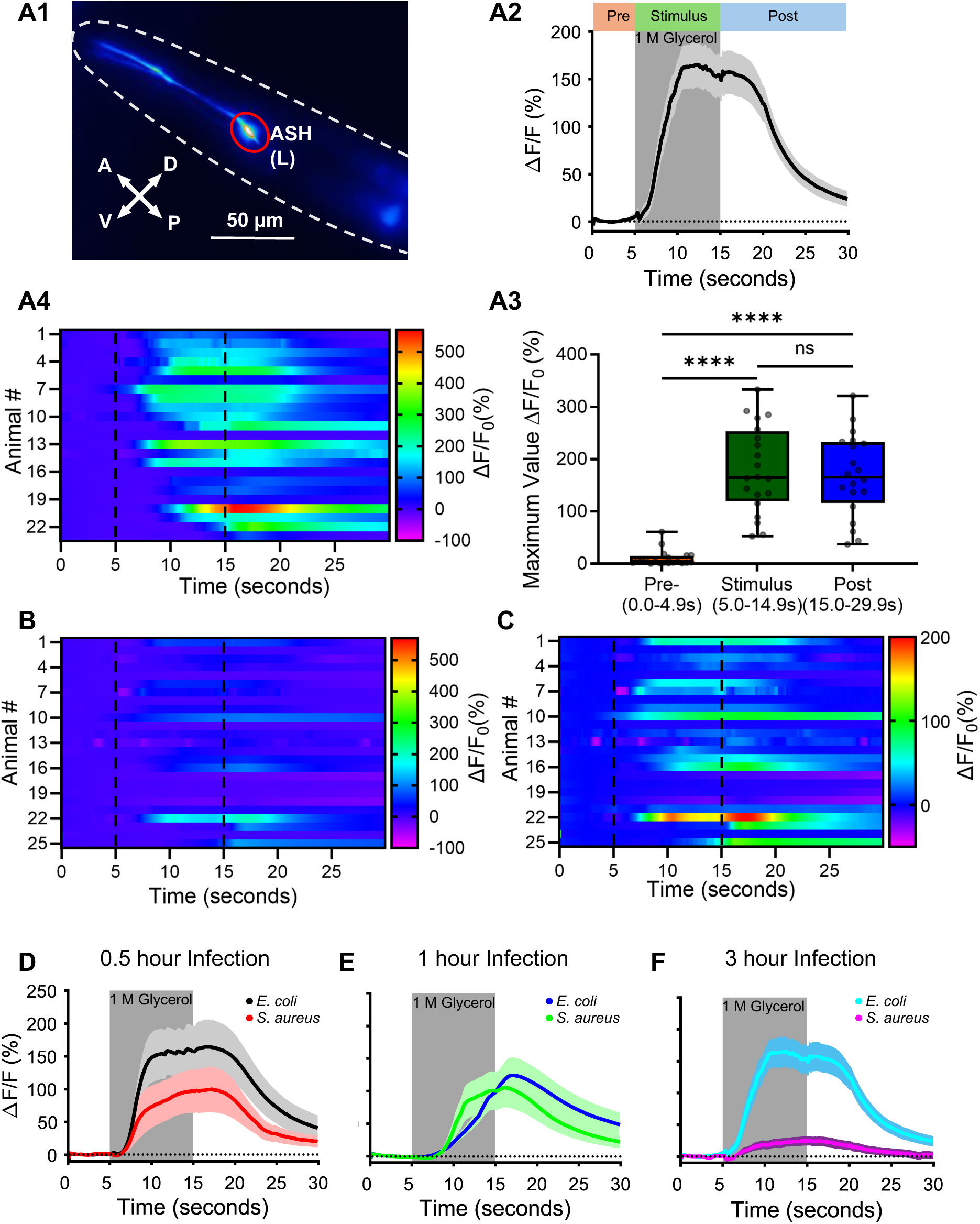
*S. aureus* infection for 3 hours results in a significant decrease in ASH neurons physiology. **A)** Annotated workflow of calcium imaging on animals fed *E. coli* for 3 hours prior to imaging with 1 M glycerol. **A1)** Animals labeled with GCaMP3.0 in ASH neuron (promoter *sra-6*), soma (outlined in red) used as region of interest. **A2)** Normalized change in fluorescence of at least 10 animals (*E. coli* fed on TSA plates for 3 hours) are averaged and plotted with SEM, grey bar indicated application of stimulus, 1 M glycerol; **A3)** the maximum change in fluorescence values of each animal is plotted at the different time points of the stimulation, Pre (0-4.9 seconds) Stimulus (5-14.9 seconds), and Post (15.0-30.0 seconds), median plotted with minimum and maximum points. (^ns^p > 0.05, ****p < 0.0001, Kruskal-Wallis test with Dunn’s multiple comparisons). **A4)** heat map plotting response of every animal (1 row = 1 animal) across time of trial, arrow indicates how one point from the maximum peak graph can be mapped to heat map. **B)** Heat map of all animals exposed to *S. aureus* for 3 hours, left same scale as **B)** **C)** Heat map of all animals exposed to *S. aureus* for 3 hours adjusted scale to display variability in response **D-F)** Percent change graphs ASH imaging of infected animals at **D)** 0.5, **E)** 1, **F)** 3 hours, stimulus of 1 M glycerol applied between 5-15 seconds.

**Supplemental Figure 2:**
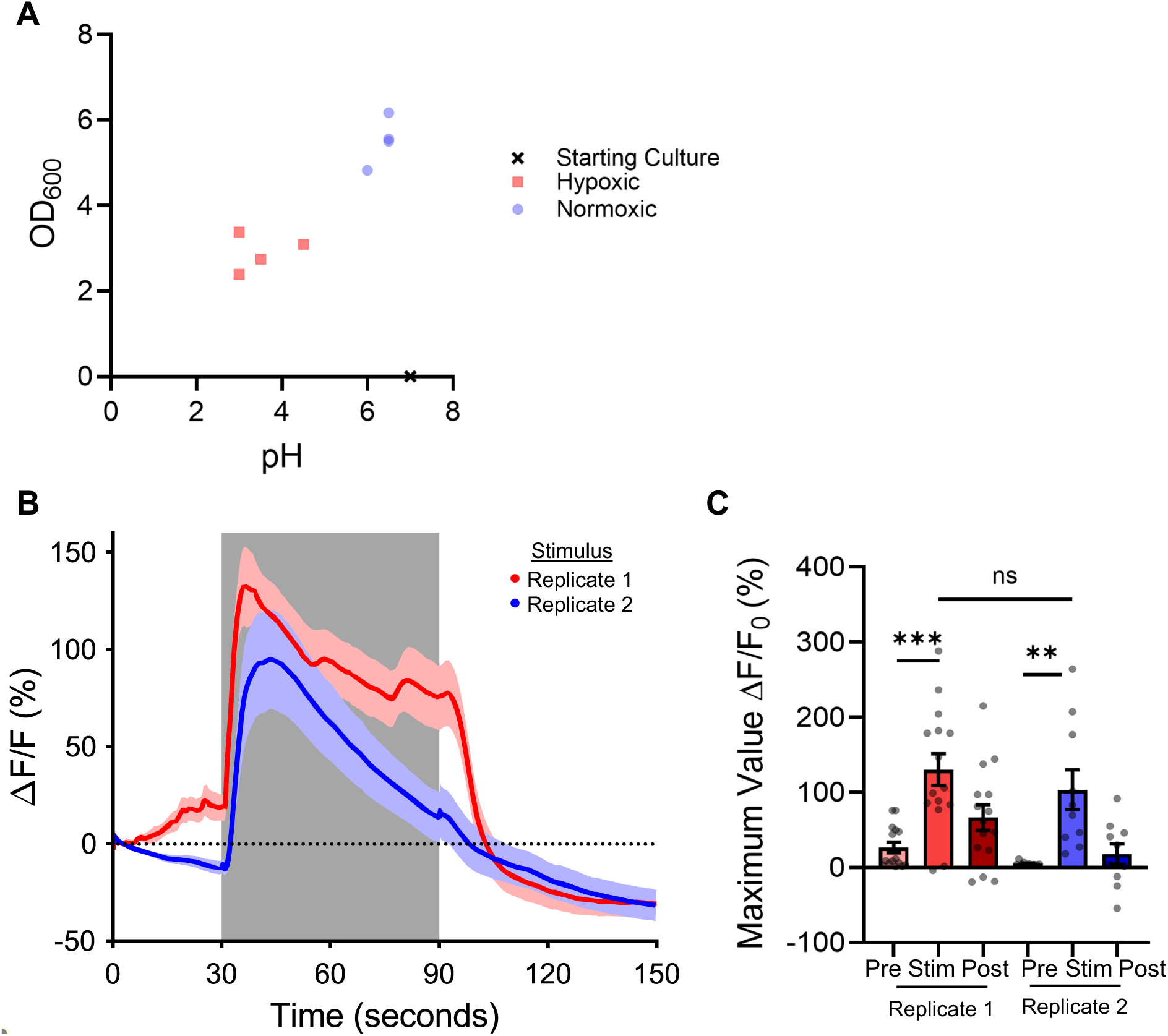
ASH activity is suppressed by stationary phase *S. aureus*. **A)** OD_600_ and pH of 5 mL *S. aureus* cultures grown under normoxic or hypoxic conditions. **B)** Response of ASH neurons to 1 M glycerol in TSB solvent, with TSB pre-exposure. Replicates performed twice throughout the course of experiments, a year apart. Trace is average response of n > 10 animals, SEM indicated with shaded region. **C)** Maximum values of fluorescence intensity of neuron with background subtracted during times of pre-stimulus exposure (0-30 seconds), stimulus exposure (30-90 seconds), and post-stimulus exposure (90-150 seconds). Each data point is the value of an individual animal. (Replicate 1 (red) parametric paired t-test ***p = 0.0003, Replicate 2 (blue) non-parametric paired Wilcoxon test **p = 0.0020, between replicated parametric Student’s unpaired two-tailed t-test ^ns^p= 0.4381. All tests with Bonferroni correction, original *p ≤ 0.05, **p ≤ 0.01, ***p ≤ 0.001, corrected *p ≤ 0.025, **p ≤ 0.005, ***p ≤ 0.0005).

**Supplemental Figure 3:**
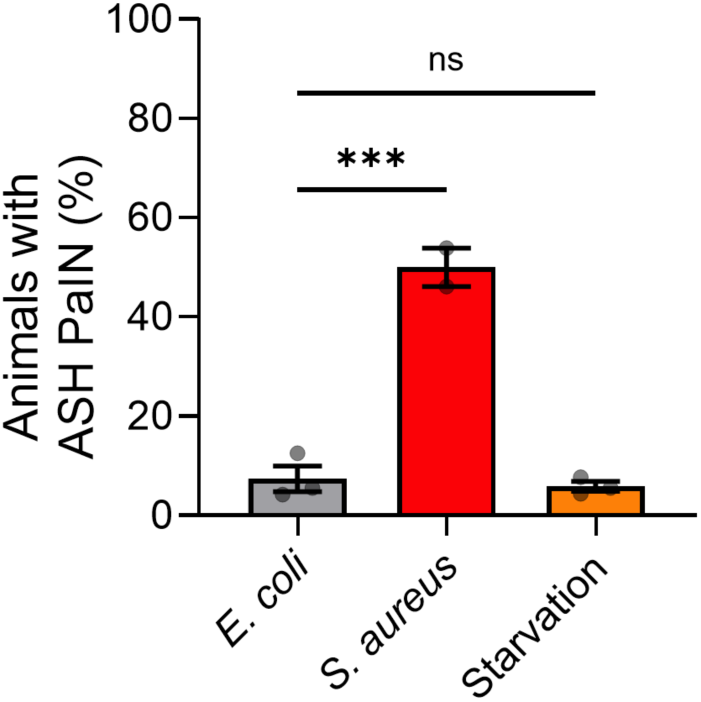
*S. aureus* infection results in Pathogen Induced Neurodegeneration (PaIN) Quantitative percent of ASH::GCaMP animals with ASH PaIN after fed *E. coli*, infected with *S. aureus*, or starved (TSA without bacteria) for 24 hours at 25 °C. Average ± SEM, three biological replicates; n = 15 - 25 animals per biological replicate. (***p = 0.0002, ^ns^p = 0.8804; Unpaired two-tail t-test with Bonferroni correction, original values ^ns^p ≥ 0.05, ***p ≤ 0.001, corrected values ^ns^p ≥ 0.025, ***p ≤ 0.0005).

## References

1. Jacobson A, Yang D, Vella M, Chiu IM. The intestinal neuro-immune axis: crosstalk between neurons, immune cells, and microbes. Mucosal Immunol. 2021;14(3):555–65.

2. Pinho-Ribeiro FA, Verri WA, Jr., Chiu IM. Nociceptor Sensory Neuron-Immune Interactions in Pain and Inflammation. Trends Immunol. 2017;38(1):5–19.

3. Gao Z, Liu Y, Zhang L, Yang Z, Lv L, Wang S, et al. Nociceptor Neurons are Involved in the Host Response to Escherichia coli Urinary Tract Infections. J Inflamm Res. 2022;15:3337–53.

4. Chiu H, Alqadah A, Chuang CF, Chang C. C. elegans as a genetic model to identify novel cellular and molecular mechanisms underlying nervous system regeneration. Cell Adh Migr. 2011;5(5):387–94.

5. Cao X, Kajino-Sakamoto R, Doss A, Aballay A. Distinct Roles of Sensory Neurons in Mediating Pathogen Avoidance and Neuropeptide-Dependent Immune Regulation. Cell Rep. 2017;21(6):1442–51.

6. Venkatesh SR, Gupta A, Singh V. Amphid sensory neurons of Caenorhabditis elegans orchestrate its survival from infection with broad classes of pathogens. Life Sci Alliance. 2023;6(8).

7. Wani KA, Goswamy D, Irazoqui JE. Nervous system control of intestinal host defense in C. elegans. Curr Opin Neurobiol. 2019;62:1–9.

8. Labed SA, Wani KA, Jagadeesan S, Hakkim A, Najibi M, Irazoqui JE. Intestinal Epithelial Wnt Signaling Mediates Acetylcholine-Triggered Host Defense against Infection. Immunity. 2018;48(5):963–78.e3.

9. Visvikis O, Ihuegbu N, Labed SA, Luhachack LG, Alves AF, Wollenberg AC, et al. Innate host defense requires TFEB-mediated transcription of cytoprotective and antimicrobial genes. Immunity. 2014;40(6):896–909.

10. Lapierre LR, De Magalhaes Filho CD, McQuary PR, Chu CC, Visvikis O, Chang JT, et al. The TFEB orthologue HLH-30 regulates autophagy and modulates longevity in Caenorhabditis elegans. Nat Commun. 2013;4:2267.

11. Irazoqui J, Troemel E, Feinbaum R, Luhachack L, Cesairliyan B, Ausubel F. Distinct Pathogenesis and Host Responses during Infection of *C. elegans* by *P. aerugunosa* and *S. aureus*. PLoS Pathogens. 2010;6(7):e1000982.

12. Liu Y, Sun J. Detection of Pathogens and Regulation of Immunity by the Caenorhabditis elegans Nervous System. mBio. 2021;12(2).

13. Vaishnava S, Yamamoto M, Severson KM, Ruhn KA, Yu X, Koren O, et al. The antibacterial lectin RegIIIgamma promotes the spatial segregation of microbiota and host in the intestine. Science (New York, NY). 2011;334(6053):255–8.

14. Treinin M, Jin Y. Cholinergic transmission in C. elegans: Functions, diversity, and maturation of ACh-activated ion channels. J Neurochem. 2021;158(6):1274–91.

15. Bargmann CI. Chemosensation in C. elegans. WormBook. 2006:1–29.

16. Hilliard MA, Apicella AJ, Kerr R, Suzuki H, Bazzicalupo P, Schafer WR. In vivo imaging of C. elegans ASH neurons: cellular response and adaptation to chemical repellents. EMBO J. 2005;24(1):63–72.

17. Hilliard MA, Bergamasco C, Arbucci S, Plasterk RH, Bazzicalupo P. Worms taste bitter: ASH neurons, QUI-1, GPA-3 and ODR-3 mediate quinine avoidance in Caenorhabditis elegans. EMBO J. 2004;23(5):1101–11.

18. Sambongi Y, Takeda K, Wakabayashi T, Ueda I, Wada Y, Futai M. Caenorhabditis elegans senses protons through amphid chemosensory neurons: proton signals elicit avoidance behavior. Neuroreport. 2000;11(10):2229–32.

19. Sassa T, Murayama T, Maruyama IN. Strongly alkaline pH avoidance mediated by ASH sensory neurons in C. elegans. Neurosci Lett. 2013;555:248–52.

20. White JG, Southgate E, Thomson JN, Brenner S. The structure of the nervous system of the nematode Caenorhabditis elegans. Philos Trans R Soc Lond B Biol Sci. 1986;314(1165):1–340.

21. Choi S, Taylor KP, Chatzigeorgiou M, Hu Z, Schafer WR, Kaplan JM. Sensory Neurons Arouse C. elegans Locomotion via Both Glutamate and Neuropeptide Release. PLoS Genet. 2015;11(7):e1005359.

22. Napolitano G, Ballabio A. TFEB at a glance. J Cell Sci. 2016;129(13):2475–81.

23. Brenner S. The genetics of Caenorhabditis elegans. Genetics. 1974;77(1):71–94.

24. Reilly DK, Lawler DE, Albrecht DR, Srinivasan J. Using an Adapted Microfluidic Olfactory Chip for the Imaging of Neuronal Activity in Response to Pheromones in Male C. Elegans Head Neurons. Journal of Visual Experimentation. 2017.

25. Chute CD, DiLoreto EM, Zhang YK, Reilly DK, Rayes D, Coyle VL, et al. Co-option of neurotransmitter signaling for inter-organismal communication in C. elegans. Nature Communications. 2019;10(1):3186.

26. Larsch J, Ventimiglia D, Bargmann CI, Albrecht DR. High-throughput imaging of neuronal activity in Caenorhabditis elegans. Proc Natl Acad Sci U S A. 2013;110(45):E4266–E73.

27. Hilliard MA, Bargmann CI, Bazzicalupo P. C. elegans responds to chemical repellents by integrating sensory inputs from the head and the tail. Curr Biol. 2002;12(9):730–4.

28. Kolter R, Siegele DA, Tormo A. The stationary phase of the bacterial life cycle. Annu Rev Microbiol. 1993;47:855–74.

29. Jenul C, Horswill AR. Regulation of Staphylococcus aureus Virulence. Microbiol Spectr. 2019;7(2).

30. Keim KC, Horswill AR. Staphylococcus aureus. Trends Microbiol. 2023;31(12):1300–1.

31. Sun JL, Zhang SK, Chen JY, Han BZ. Metabolic profiling of Staphylococcus aureus cultivated under aerobic and anaerobic conditions with (1)H NMR-based nontargeted analysis. Can J Microbiol. 2012;58(6):709–18.

32. Fuchs S, Pané-Farré J, Kohler C, Hecker M, Engelmann S. Anaerobic gene expression in Staphylococcus aureus. J Bacteriol. 2007;189(11):4275–89.

33. Kohler C, Wolff S, Albrecht D, Fuchs S, Becher D, Büttner K, et al. Proteome analyses of Staphylococcus aureus in growing and non-growing cells: a physiological approach. Int J Med Microbiol. 2005;295(8):547–65.

34. Bae YK, Barr MM. Sensory roles of neuronal cilia: cilia development, morphogenesis, and function in C. elegans. Front Biosci. 2008;13:5959–74.

35. Dirksen P, Assié A, Zimmermann J, Zhang F, Tietje AM, Marsh SA, et al. CeMbio - The Caenorhabditis elegans Microbiome Resource. G3 (Bethesda). 2020;10(9):3025–39.

36. Gonzalez X, Irazoqui JE. Distinct members of the Caenorhabditis elegans CeMbio reference microbiota exert cryptic virulence that is masked by host defense. Mol Microbiol. 2024;122(3):387–402.

37. Meng L, Mulcahy B, Cook SJ, Neubauer M, Wan A, Jin Y, et al. The Cell Death Pathway Regulates Synapse Elimination through Cleavage of Gelsolin in Caenorhabditis elegans Neurons. Cell Rep. 2015;11(11):1737–48.

38. Crook M, Upadhyay A, Hanna-Rose W. Necrosis in C. elegans. Methods Mol Biol. 2013;1004:171–82.

39. Tschuck J, Padmanabhan Nair V, Galhoz A, Zaratiegui C, Tai HM, Ciceri G, et al. Suppression of ferroptosis by vitamin A or radical-trapping antioxidants is essential for neuronal development. Nat Commun. 2024;15(1):7611.

40. Chen X, Wang Y, Chen YZ, Harry BL, Nakagawa A, Lee ES, et al. Regulation of CED-3 caspase localization and activation by C. elegans nuclear-membrane protein NPP-14. Nat Struct Mol Biol. 2016;23(11):958–64.

41. Xue D, Horvitz HR. Caenorhabditis elegans CED-9 protein is a bifunctional cell-death inhibitor. Nature. 1997;390(6657):305–8.

42. Syntichaki P, Xu K, Driscoll M, Tavernarakis N. Specific aspartyl and calpain proteases are required for neurodegeneration in C. elegans. Nature. 2002;419(6910):939–44.

43. Xu K, Tavernarakis N, Driscoll M. Necrotic cell death in C. elegans requires the function of calreticulin and regulators of Ca(2+) release from the endoplasmic reticulum. Neuron. 2001;31(6):957–71.

44. Walker DS, Gower NJ, Ly S, Bradley GL, Baylis HA. Regulated disruption of inositol 1,4,5-trisphosphate signaling in Caenorhabditis elegans reveals new functions in feeding and embryogenesis. Mol Biol Cell. 2002;13(4):1329–37.

45. Alvarez J, Alvarez-Illera P, García-Casas P, Fonteriz RI, Montero M. The Role of Ca(2+) Signaling in Aging and Neurodegeneration: Insights from Caenorhabditis elegans Models. Cells. 2020;9(1).

46. Dixon SJ, Stockwell BR. The Hallmarks of Ferroptosis. Annual Review of Cancer Biology. 2019;3(Volume 3, 2019):35–54.

47. Dixon SJ, Lemberg KM, Lamprecht MR, Skouta R, Zaitsev EM, Gleason CE, et al. Ferroptosis: an iron-dependent form of nonapoptotic cell death. Cell. 2012;149(5):1060–72.

48. Perez MA, Magtanong L, Dixon SJ, Watts JL. Dietary Lipids Induce Ferroptosis in Caenorhabditiselegans and Human Cancer Cells. Dev Cell. 2020;54(4):447–54.e4.

49. Hillyard SL, German JB. Quantitative lipid analysis and life span of the fat-3 mutant of Caenorhabditis elegans. J Agric Food Chem. 2009;57(8):3389–96.

50. Cecconi F, Levine B. The role of autophagy in mammalian development: cell makeover rather than cell death. Dev Cell. 2008;15(3):344–57.

51. Suomi F, McWilliams TG. Autophagy in the mammalian nervous system: a primer for neuroscientists. Neuronal Signal. 2019;3(3):Ns20180134.

52. Wani KA, Goswamy D, Taubert S, Ratnappan R, Ghazi A, Irazoqui JE. NHR-49/PPAR-α and HLH-30/TFEB cooperate for C. elegans host defense via a flavin-containing monooxygenase. eLife. 2021;10.

53. Minnerly J, Zhang J, Parker T, Kaul T, Jia K. The cell non-autonomous function of ATG-18 is essential for neuroendocrine regulation of Caenorhabditis elegans lifespan. PLoS Genet. 2017;13(5):e1006764.

54. Wu F, Li Y, Wang F, Noda NN, Zhang H. Differential function of the two Atg4 homologues in the aggrephagy pathway in Caenorhabditis elegans. J Biol Chem. 2012;287(35):29457–67.

55. Sambongi Y, Nagae T, Liu Y, Yoshimizu T, Takeda K, Wada Y, et al. Sensing of cadmium and copper ions by externally exposed ADL, ASE, and ASH neurons elicits avoidance response in Caenorhabditis elegans. Neuroreport. 1999;10(4):753–7.

56. Bargmann CI, Mori I. Chemotaxis and Thermotaxis. In: Riddle DL, Blumenthal T, Meyer BJ, Priess JR, editors. C elegans II. Cold Spring Harbor (NY): Cold Spring Harbor Laboratory Press Copyright © 1997, Cold Spring Harbor Laboratory Press.; 1997.

57. McLachlan IG, Kramer TS, Dua M, DiLoreto EM, Gomes MA, Dag U, et al. Diverse states and stimuli tune olfactory receptor expression levels to modulate food-seeking behavior. eLife. 2022;11.

58. Prakash D, Ms A, Radhika B, Venkatesan R, Chalasani SH, Singh V. 1-Undecene from Pseudomonas aeruginosa is an olfactory signal for flight-or-fight response in Caenorhabditis elegans. EMBO J. 2021;40(13):e106938.

59. Tam K, Torres VJ. Staphylococcus aureus Secreted Toxins and Extracellular Enzymes. Microbiol Spectr. 2019;7(2).

60. Harper L, Balasubramanian D, Ohneck EA, Sause WE, Chapman J, Mejia-Sosa B, et al. Staphylococcus aureus Responds to the Central Metabolite Pyruvate To Regulate Virulence. mBio. 2018;9(1).

61. Seilie ES, Bubeck Wardenburg J. Staphylococcus aureus pore-forming toxins: The interface of pathogen and host complexity. Semin Cell Dev Biol. 2017;72:101–16.

62. Pinho-Ribeiro FA, Baddal B, Haarsma R, O’Seaghdha M, Yang NJ, Blake KJ, et al. Blocking Neuronal Signaling to Immune Cells Treats Streptococcal Invasive Infection. Cell. 2018;173(5):1083–97.e22.

63. Wu Q, Cao X, Yan D, Wang D, Aballay A. Genetic Screen Reveals Link between the Maternal Effect Sterile Gene mes-1 and Pseudomonas aeruginosa-induced Neurodegeneration in Caenorhabditis elegans. J Biol Chem. 2015;290(49):29231–9.

64. Lezi E, Zhou T, Koh S, Chuang M, Sharma R, Pujol N, et al. An Antimicrobial Peptide and Its Neuronal Receptor Regulate Dendrite Degeneration in Aging and Infection. Neuron. 2018;97(1):125–38.e5.

65. Rawson RL, Yam L, Weimer RM, Bend EG, Hartwieg E, Horvitz HR, et al. Axons degenerate in the absence of mitochondria in C. elegans. Curr Biol. 2014;24(7):760–5.

66. Kaur S, Aballay A. G-Protein-Coupled Receptor SRBC-48 Protects against Dendrite Degeneration and Reduced Longevity Due to Infection. Cell Rep. 2020;31(7):107662.

67. Kaur S, Sang Y, Aballay A. Myotubularin-related protein protects against neuronal degeneration mediated by oxidative stress or infection. J Biol Chem. 2022;298(3):101614.

68. Cook SJ, Jarrell TA, Brittin CA, Wang Y, Bloniarz AE, Yakovlev MA, et al. Whole-animal connectomes of both Caenorhabditis elegans sexes. Nature. 2019;571(7763):63–71.

69. Settembre C, Di Malta C, Polito VA, Garcia Arencibia M, Vetrini F, Erdin S, et al. TFEB links autophagy to lysosomal biogenesis. Science (New York, NY). 2011;332(6036):1429–33.

70. Yan C, Liu J, Gao J, Sun Y, Zhang L, Song H, et al. IRE1 promotes neurodegeneration through autophagy-dependent neuron death in the Drosophila model of Parkinson’s disease. Cell Death & Disease. 2019;10(11):800.

71. Palmer JE, Wilson N, Son SM, Obrocki P, Wrobel L, Rob M, et al. Autophagy, aging, and age-related neurodegeneration. Neuron. 2025;113(1):29–48.

72. Guo F, Liu X, Cai H, Le W. Autophagy in neurodegenerative diseases: pathogenesis and therapy. Brain Pathol. 2018;28(1):3–13.

73. Nixon RA. Autophagy-lysosomal-associated neuronal death in neurodegenerative disease. Acta Neuropathol. 2024;148(1):42.

74. Lotz SK, Blackhurst BM, Reagin KL, Funk KE. Microbial Infections Are a Risk Factor for Neurodegenerative Diseases. Front Cell Neurosci. 2021;15:691136.

75. Hammarlund M, Hobert O, Miller DM, 3rd, Sestan N. The CeNGEN Project: The Complete Gene Expression Map of an Entire Nervous System. Neuron. 2018;99(3):430–3.

76. Takla M, Keshri S, Rubinsztein DC. The post-translational regulation of transcription factor EB (TFEB) in health and disease. EMBO Rep. 2023;24(11):e57574.

77. Jiao F, Zhou B, Meng L. The regulatory mechanism and therapeutic potential of transcription factor EB in neurodegenerative diseases. CNS Neurosci Ther. 2023;29(1):37–59.

